# Single-molecule displacement mapping unveils nanoscale heterogeneities and charge effects in intracellular diffusivity

**DOI:** 10.1101/559484

**Authors:** Limin Xiang, Kun Chen, Rui Yan, Wan Li, Ke Xu

**Affiliations:** Department of Chemistry, University of California, Berkeley, CA 94720, USA; Chan Zuckerberg Biohub, San Francisco, CA 94158, USA

**Keywords:** Intracellular diffusion, super-resolution microscopy, single-molecule spectroscopy, macromolecular crowding, protein net charge

## Abstract

Intracellular diffusion underlies vital cellular processes. However, it remains difficult to elucidate how an average-sized protein diffuses inside the cell with good spatial resolution and sensitivity. Here we introduce single-molecule displacement/diffusivity mapping (SM*d*M), a super-resolution strategy that enables the nanometer-scale mapping of intracellular diffusivity through the local statistics of instantaneous displacements of freely diffusing single molecules. We thus show that diffusion in the mammalian cytoplasm and nucleus to both be spatially heterogeneous at the nanoscale, and that such variations in local diffusivity correlate well with the ultrastructure of the actin cytoskeleton and the chromosome, respectively. Moreover, we identify the net charge of the diffuser as a key determinant of diffusion rate: intriguingly, the possession of positive, but not negative, net charges significantly impedes diffusion, and the exact degree of slowdown is determined by the specific subcellular environments. We thus open a new door to understanding the physical rules that govern the intracellular interactions between biomolecules at the nanoscale.

## INTRODUCTION

The magic of life occurs when the right molecules meet. Whereas active transport provides an organized, yet costly means to move things around inside the eukaryotic cell, passive diffusion offers a mechanism for molecules to mix “for free”. It, however, remains difficult to map out how an average-sized protein diffuses in the live cell with good spatial resolution and sensitivity. Does intracellular diffusivity contain structures at the nanoscale, and if so, how are they modulated by the local intracellular structures and microenvironments, as well as by the properties of the diffuser itself?

Although environment-sensitive probes have been developed to directly visualize intracellular parameters and processes as viscosity, macromolecular crowding, and protein-folding dynamics (Boersma et al., 2015; Ebbinghaus et al., 2010; Kuimova et al., 2009; Rivas and Minton, 2016; Wirth and Gruebele, 2013; Yang et al., 2014), they do not address diffusivity. Photobleaching and photoactivation-based techniques (Ishikawa-Ankerhold et al., 2012; Lippincott-Schwartz et al., 2001) enable single-location diffusion measurements, but are unamicable to spatial mapping. Fluorescence correlation spectroscopy (FCS) and related methods (Digman and Gratton, 2011; Machan and Wohland, 2014; Ries and Schwille, 2012) infer diffusivity from spatiotemporal fluctuations in intensity, but are sensitive to experimental conditions (Enderlein et al., 2005; Ries and Schwille, 2012) and achieve limited resolution and sensitivity in live cells.

Single-molecule tracking has been highly successful for membrane- and chromosome-bound molecules and for molecules diffusing inside the confined volumes of bacteria (Cognet et al., 2014; Elf and Barkefors, 2019; Kusumi et al., 2014; Manley et al., 2008; Manzo and Garcia-Parajo, 2015). However, it remains challenging to apply single-molecule tracking to unbound molecules freely diffusing inside the eukaryotic cell. To record a reasonably large area, modern high-sensitivity cameras often limit time resolution to ~10 ms (~100 frames per second). For an average-sized protein with an intracellular diffusion coefficient *D* of ~20-30 μm^2^/s (Lippincott-Schwartz et al., 2001; Milo and Phillips, 2016), this frame time results in ~700 nm of diffusion in each dimension, hence severe motion-blur. Although stroboscopic illumination overcomes motion-blur (Elf et al., 2007; English et al., 2011), tracking between frames remains difficult for the eukaryotic cell: with ~700 nm axial displacement, a molecule initially in focus readily diffuses out of the focal range (~±400 nm for a high-NA objective) in the subsequent frame (see below), an issue not encountered in bacteria for their very small dimensions.

We here develop a strategy to first determine the nanoscale displacements of freely diffusing single molecules in short (~1 ms) time windows through the application of a pair of closely timed excitation pulses. By repeating such pulse pairs for ~10^4^ times and locally accumulating the resultant single-molecule *displacements*, we next construct super-resolution maps of diffusion rate, and hence uncover nanoscale diffusivity heterogeneities in live mammalian cells. We name this strategy single-molecule displacement/diffusivity mapping (SM*d*M), a tribute to single-molecule localization microscopy (SMLM), which generates super-resolution images by accumulating single-molecule *localizations* (Betzig et al., 2006; Hess et al., 2006; Rust et al., 2006).

## RESULTS

### SM*d*M enables super-resolution mapping of intracellular diffusivity via local statistics of the instantaneous displacements of freely diffusing single molecules

We first expressed free mEos3.2, a photoswitchable, monomeric fluorescent protein (FP) commonly used in SMLM (Zhang et al., 2012), in the cytoplasm of mammalian cells. Along with a short cloning-site sequence, the expressed protein (mEos3.2-C1; Table S1) contained 252 amino acids (AA) (~28 kDa), close to the medium size of human proteins [248 AA by abundance (Milo and Phillips, 2016)]. As with typical SMLM experiments, we illuminated several micrometers into the coverslip-adhered live cells with a 561 nm excitation laser, and used a weak 405 nm laser to photoswitch a small fraction of the expressed mEos3.2 molecules to the 561 nm-excitable state, hence a means to control the amount of fluorescent single molecules in the view (Betzig et al., 2006; Manley et al., 2008). As expected, at a 109 Hz framerate (camera frame time *T* = 9.16 ms), freely diffusing single mEos3.2 molecules appeared blurry (Figure 1A). The application of stroboscopic illumination (Elf et al., 2007; English et al., 2011), in which excitation pulses *τ* = 500 μs in duration were synchronized to the center of each camera frame, provided clear single-molecule images (Figure 1B). However, in the succeeding frame, after the frame time of *T* = 9.16 ms, molecules detected in the first frame already diffused out of the focal range and so could not be tracked (Figure 1B).

**Figure 1.**
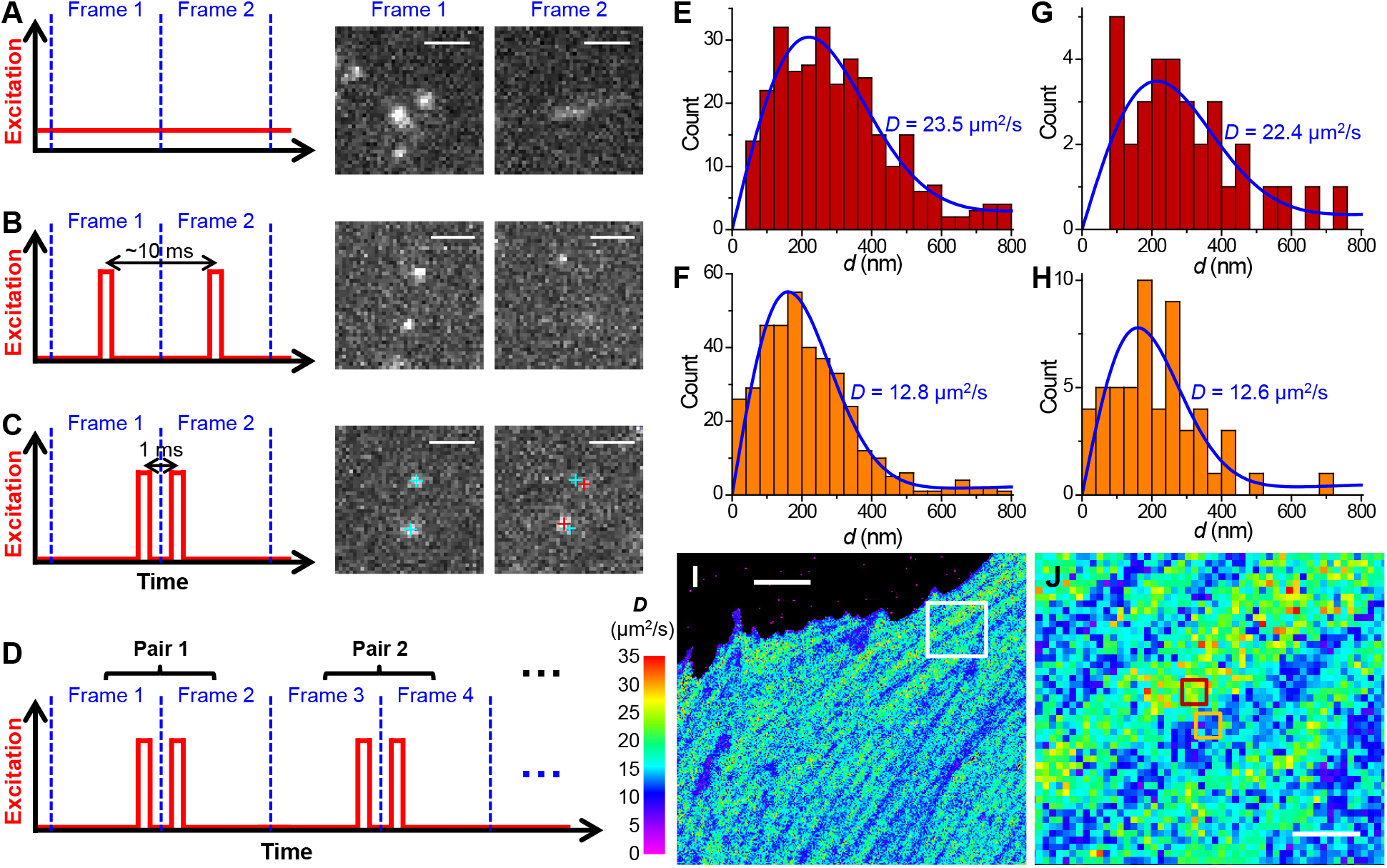
Super-resolution displacement mapping of single mEos3.2 FP molecules freely diffusing in the cytoplasm of live mammalian cells. (A) Conventional imaging with continuous laser illumination and a recording framerate of 109 Hz. (B) Stroboscopic illumination, with excitation pulses *τ* = 500 μs in duration synchronized to the center of each camera frame. (C) Placing two excitation pulses towards the end of the first frame and the beginning of the second frame, respectively, so that the center-to-center time separation between the two recorded images is reduced to 1 ms. Cyan and red crosses mark the super-localized positions of two detected molecules in Frame 1 and Frame 2, respectively. (D) Such paired frames are repeated ~10^4^ times to enable statistics. (E,F) Distribution of *d* for two 300×300 nm^2^ areas [red and orange boxes in (J)]. (G,H) Distribution of *d* for two 100×100 nm^2^ areas at the centers of the areas for (E,F), respectively. Blue lines in (E-H) are MLE results using eqn. 2 in Methods, with corresponding *D* values labeled in each figure. (I,J) Map of intracellular diffusivity constructed through MLE of the *d* distribution in every 100×100 nm^2^ spatial bin. (J) is a zoom-in of the white box in (I). Scale bars: 2 μm (A-C), 5 μm (I), 1 μm (J). See also Figure S1.

To overcome this issue, we reduced the temporal separation between the pair of captured images by placing two excitation pulses towards the end of the first frame and the beginning of the second frame, respectively (Figure 1C). Thus, at a ∆*t* = 1 ms center-to-center separation between the two pulses, molecules being detected in the first frame (due to the first pulse) had only traveled moderately (to stay within focus) at the time of the second pulse (captured in the second frame) (Figure 1C). Comparing the super-localized positions of the molecules in the two frames thus yielded their nanoscale displacements (*d*) in the Δ*t* = 1 ms time window.

We next repeated recording ~10^4^ pairs of frames to enable statistics (Figure 1D). The temporal proximity of the paired excitation pulses (Δ*t*) left ample time between the unpaired pulses (2*T*−Δ*t*) for different molecules to diffuse into the focal range as independent reporters of local diffusivity. The resultant, accumulated *d* values were spatially binned to evaluate local *D*. At a 300×300 nm^2^ bin size (Figures 1E and 1F), the distribution of *d* in each bin was well fitted by a modified two-dimensional random-walk model (Methods) through maximum likelihood estimation (MLE). Reducing the bin size to 100×100 nm^2^ led to increased statistical uncertainties for each bin, but MLE still yielded reasonable results (Figures 1G and 1H). We further demonstrated the robustness of our fitting model for high single-molecule density (Figure S1). Color-plotting the *D* values obtained by individually performing MLE for each 100×100 nm^2^ spatial bin thus rendered a super-resolution map of local *D* across the full view (Figures 1I and 1J).

### Diffusivity in the mammalian cytoplasm is spatially heterogeneous at the nanoscale due to the actin cytoskeleton

For mEos3.2 molecules freely diffusing in the cytoplasm of live mammalian cells, we observed typical *D* of 20-25 μm^2^/s for the high-*D* regions (Figures 1I, 1J, 2A, 2C, S1, and S2), comparable to previous, spatially unresolved results of FPs (Lippincott-Schwartz et al., 2001; Milo and Phillips, 2016). Treating the cells with a 2× hyperosmotic medium led to substantially reduced *D* down to ~8 μm^2^/s for the high-*D* regions (Figure 2B), consistent with increased macromolecular crowding owning to water loss (Boersma et al., 2015; Swaminathan et al., 1997).

**Figure 2.**
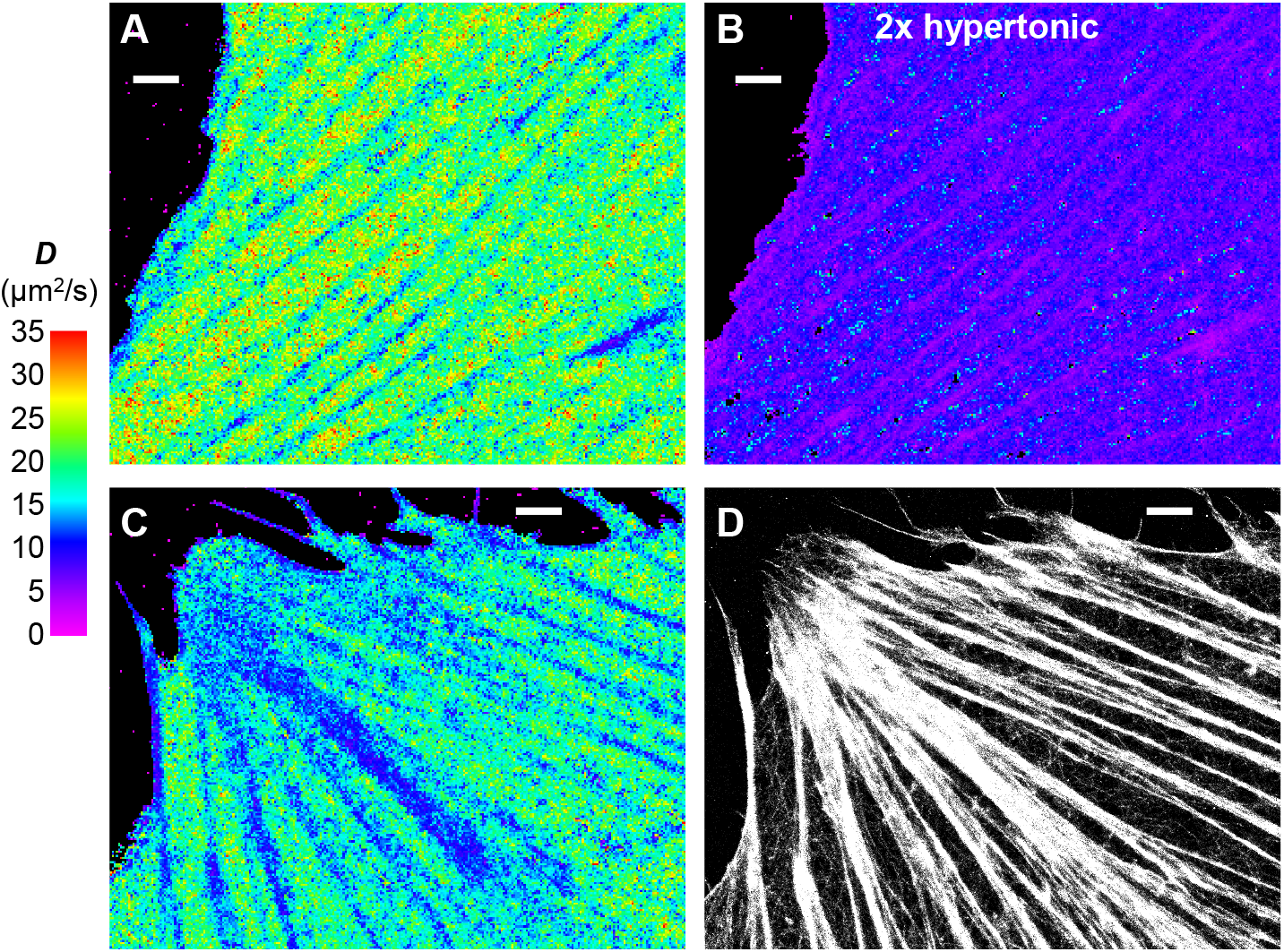
Diffusivity in the mammalian cytoplasm is spatially heterogeneous at the nanoscale due to the actin cytoskeleton. (A) SM*d*M diffusivity map of free mEos3.2 molecules in the cytoplasm of a live PtK2 cell. (B) The same cell in a 2× hyperosmotic medium. (C,D) Correlated SM*d*M diffusivity map of mEos3.2 in another live PtK2 cell (C), vs. SMLM image of Alexa Fluor 647 phalloidin-labeled actin in the fixed cell (D). Scale bars in all panels: 2 μm. See also Figure S2.

Meanwhile, our ability to map local *D* throughout the cell revealed substantial diffusivity heterogeneities at the nanoscale. For the flat, spread parts of cells, SM*d*M *D* maps often showed continuous, linear features characterized by markedly reduced *D* values down to ~10 μm^2^/s (Figures 1I, 2A, 2C, and S2). The distinct linear structures are reminiscent of the actin cytoskeleton, which often form linear bundles as stress fibers. Indeed, SMLM of the phalloidin-labeled fixed cell (Xu et al., 2012) showed good correspondences between actin bundles and the SM*d*M-revealed low-*D* regions in the live cell (Figures 2C, 2D, and S2).

### Diffusivity in the mammalian nucleus is spatially heterogeneous at the nanoscale due to the nucleolus and the chromatin

We next examined diffusion in the nucleus. By setting the focal plane a few micrometers into the cell, we imaged at the central depths of the nuclei. SM*d*M (Figures 3A and S3) yielded *D* of ~20 μm^2^/s for the highest-*D* regions of the nucleus (red arrows), consistent with the view that the nucleosol and cytosol share similar diffusion properties (Seksek et al., 1997). Meanwhile, micrometer-sized globule structures were noted, where the local *D* dropped substantially to ~6 μm^2^/s (white asterisk in Figure 3A). The globule shape is reminiscent of the nucleolus, a subnuclear compartment for ribosome biogenesis (Boisvert et al., 2007). Our bright-field transmission images supported this assignment (Figure S3). The observed, much-reduced *D* in the nucleolus is consistent with its high crowdedness of proteins and nucleic acids (Boisvert et al., 2007).

**Figure 3.**
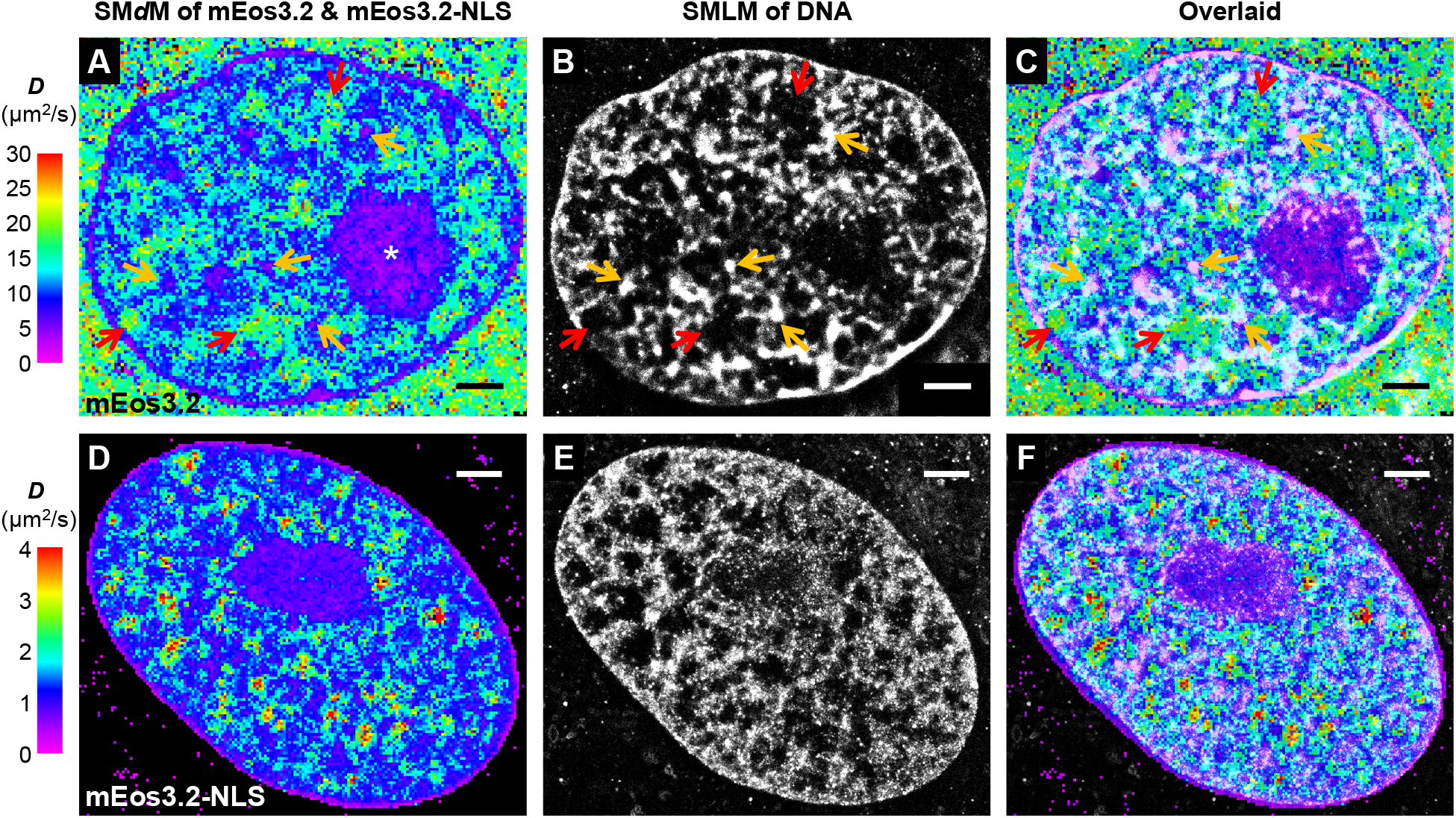
Diffusivity in the mammalian nucleus is spatially heterogeneous at the nanoscale due to the nucleolus and the chromatin. (A) SM*d*M diffusivity map of free mEos3.2 at the central depth of the nucleus of a live PtK2 cell. (B) SMLM image of the fixed cell using the DNA stain NucSpot Live 650. (C) Overlay of (A) and (B). (D) SM*d*M diffusivity map of mEos3.2-NLS in the nucleus of a live PtK2 cell. (E) SMLM image of NucSpot-stained DNA of the fixed cell. (F) Overlay of (D) and (E). Scale bars in all panels: 2 μm. See also Figure S3.

Close examination of the SM*d*M data further revealed semi-structured, fractal-like nanoscale features of lowered *D* (~10 μm^2^/s), which sporadically evolved into ~200 nm sized foci of very low *D* of ~6 μm^2^/s (orange arrows in Figure 3A). To examine if these features were related to the chromatin ultrastructure, we performed SMLM on the fixed cell with a DNA stain. This showed the coexistence of dense chromatin fibers and nanoscale voids (Figure 3C), consistent with recent super-resolution observations (Benke and Manley, 2012; Ricci et al., 2015; Szczurek et al., 2017). Remarkably, a strong correlation was found between the SM*d*M map of mEos3.2 and the SMLM image of DNA: the highest *D* values were consistently observed for regions devoid of DNA (red arrows in Figures 3A-3C), whereas the low *D* regions corresponded to DNA structures, with the slowest foci often corresponding to clusters of high local DNA density (orange arrows), a structure indicative of densely packed structures as the heterochromatin. See Figure S3 for additional examples: the spatial patterns of diffusivity correlated well with diverse chromatin ultrastructures. These results implicate chromatin crowding as a major impediment to intranuclear diffusion.

### An unexpected, substantial slowdown in diffusion by the nuclear localization sequence

For specific visualization of diffusion inside the nucleus, we further added a nuclear localization sequence (NLS) (Marfori et al., 2011) to mEos3.2 (Table S1). Unexpectedly, although SM*d*M maps of mEos3.2-NLS again correlated well with the SMLM-resolved DNA (Figures 3D-3F), the actual *D* values dropped by one order of magnitude (Figure 3D). As this big drop is unlikely due to the small added size (262 vs. 252 AAs), we questioned what alternative factors could have dominated the diffusion behavior, and noticed the strong positive charge of NLS: Under the physiological pH of 7.4, our original mEos3.2 expression had a net charge of +2. With the NLS, the net charge became +15 (Table S1).

### The possession of positive, but not negative, net charges is a key determinant of diffusivity in the mammalian cell

To test the possible effect of protein charge on intracellular diffusion, we started by adding short, consecutive Asp/Glu and Arg/Lys sequences to the C-terminus of the expressed mEos3.2 protein, yielding net charges of −14, −7, 0, +7, and +14 (Table S1). SM*d*M showed a surprising trend: For all subcellular environments, the two negatively charged (−14, −7) species (Figures 4A and 4B) both yielded *D* comparable to that of the neutral (0 charges) species (Figure 4C), but slightly higher than that of the original mEos3.2-C1 (+2 charges) (Figures 2, 3, and 4F). For the more positively charged protein (+7), however, markedly reduced *D*, down to half of that of the negative and neutral species, was found across all subcellular environments (Figures 4D and 4F). Meanwhile, extremely slow diffusion was found for the +14 charged protein (Figure 4E; note the reduced color scale for diffusion rate): Curiously, as *D* dropped to ~0.5 μm^2^/s in the cytoplasm, notably higher values of up to ~3 μm^2^/s were retained inside the nucleus, comparable to what we initially noticed for mEos3.2-NLS (+15 charges; Figure 3D).

**Figure 4.**
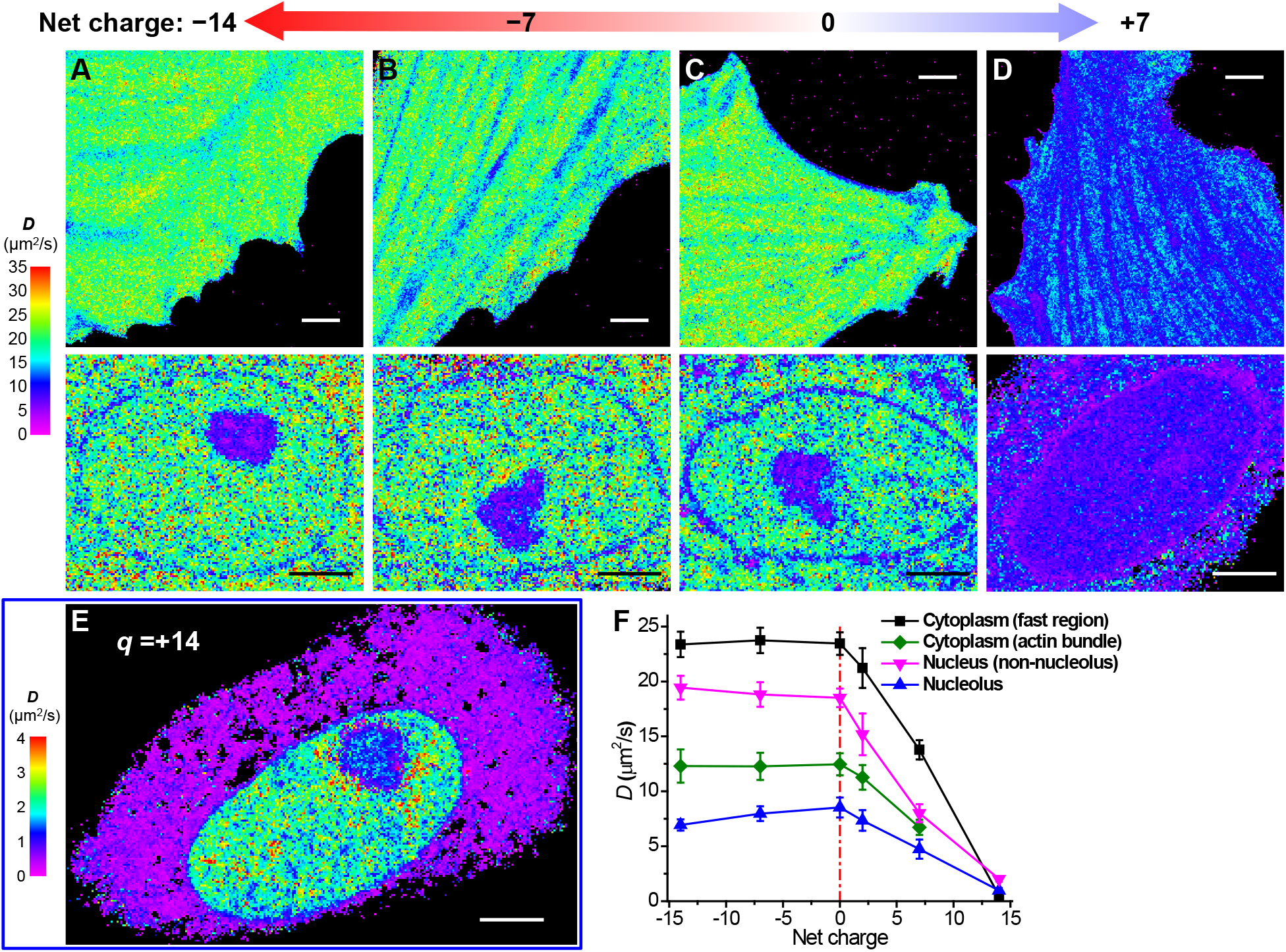
The possession of positive, but not negative, net charges is a key determinant of diffusivity in the mammalian cell. (A-D) SM*d*M diffusivity maps of mEos3.2 constructs of −14 (A), −7 (B), 0 (C), and +7 (D) net charges expressed in live PtK2 cells. The top and bottom panels show representative results in the spread parts of cells and the nuclei, respectively. (E) SM*d*M diffusivity map of +14 charged mEos3.2 in a live PtK2 cell, on a substantially reduced *D* scale. (F) Mean *D* values for the above differently charged proteins in different subcellular environments. For cytoplasm, averaged *D* is presented for fast regions with no apparent slowdown due to the actin cytoskeleton (black), as well as for the actin-bundle regions (green). The nucleus data are simply divided into nucleolus (blue) and non-nucleolus (magenta) regions. Error bars: standard deviations between individual cells (*n* > 6 cells for each data point). Scale bars: 4 μm (A-E).

To elucidate whether the above-observed diffusion slowdown of the positively charged proteins was specific to motifs of consecutive Arg/Lys, which might bind to importins (Marfori et al., 2011) or DNA (Xiong and Blainey, 2016), we further examined two other proteins of +7 net charge, one containing 7 sparsely distributed Arg/Lys in a 21 AA sequence at the C-terminus of mEos3.2 (+7b; Table S1), and another with modifications to the mEos3.2 sequence at 3 well-separated locations (+7c; Table S1). Notably, similar degrees of diffusion slowdown (vs. the neutral protein, Figures 4C and 5A) were found for both proteins (Figures 5C and 5D) when compared to the original one with consecutive Arg/Lys (+7a; Figures 4D and 5B), with similar trends observed across different subcellular environments (Figure 5E). This result indicates that it was the effect of net charge, rather than specific interactions due to particular protein sequences, that drove the different diffusion behaviors.

**Figure 5.**
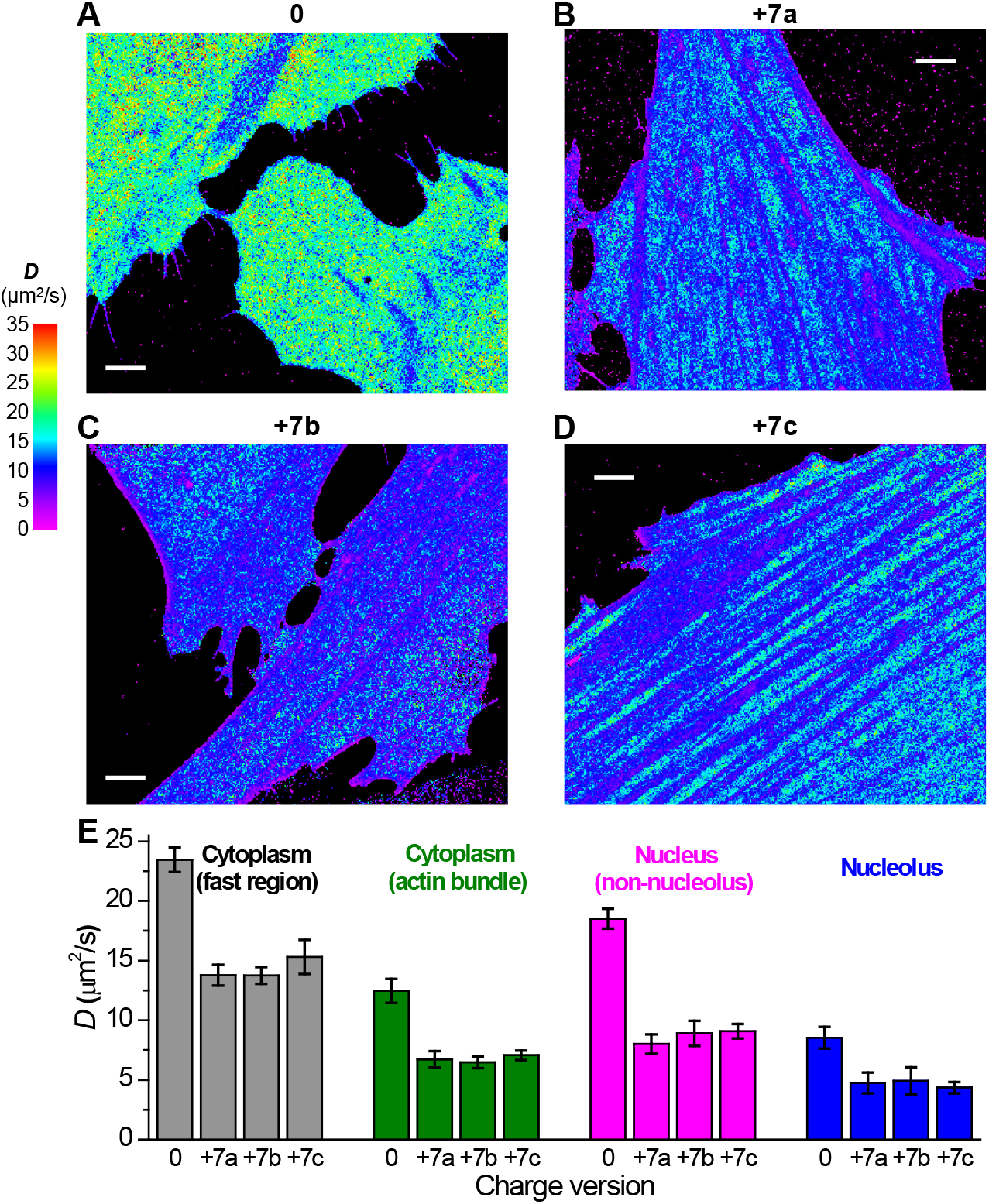
Results on different +7 charged species point to the effect of positive net charge, rather than specific sequences, in diffusion slowdown. (A) Another example of SM*d*M diffusivity map of 0 charged mEos3.2 in live PtK2 cells. (B-D) Representative SM*d*M diffusivity maps for three different versions of +7 charged FPs in live PtK2 cells. (B) +7a: consecutive Arg/Lys at the C-terminus of mEos3.2. (C) +7b: 7 Arg/Lys distributed over 21 AAs at the C-terminus of mEos3.2. (D) +7c: modifications to the mEos3.2 sequence at 3 well-separated locations. See also Table S1. (E) Comparison of the mean *D* values for the above proteins in different subcellular environments. Error bars: standard deviations between individual cells (n > 6 cells for each data point). Scale bars: 4 μm (A-D).

## DISCUSSION

Intracellular diffusion underlies fundamental processes of the cell. However, it has been a big challenge to elucidate how an average-sized, unbound protein diffuses intracellularly with reasonable spatial resolution and sensitivity. In particular, whereas single-molecule tracking has been powerful in examining the diffusion behavior of membrane- and chromosome-bound molecules and for volume-confined systems like bacteria, its need to follow each molecule over consecutive frames makes its application to the fast, free diffusion inside eukaryotic cells impractical.

SM*d*M eliminated the need to track each molecule over multiple frames, and flipped the question to evaluate, for each fixed location, how different single molecules travel locally. Thus, by devising a paired excitation scheme to record, in 1 ms time windows, the nanoscale displacements of different molecules that stochastically entered the focal plane, super-resolution diffusivity maps were generated for freely diffusing molecules. Consequently, we unveiled nanoscale diffusivity heterogeneities in both the mammalian cytoplasm and nucleus.

For the cytoplasm, we observed local diffusion slowdown that corresponded to the actin cytoskeleton. Previous work has suggested that the actin cytoskeleton impedes intracellular diffusion at the whole-cell level (Baum et al., 2014; Potma et al., 2001). Meanwhile, imaging with viscosity-sensing dyes detects no distinct intracellular structures (Kuimova et al., 2009; Yang et al., 2014). SM*d*M resolved nanoscale heterogeneity in *D*, and directly linked substantial decreases in *D* to the local actin ultrastructure. Interestingly, a protein-folding sensor has shown linear intracellular features of locally elevated melting temperature (Ebbinghaus et al., 2010; Wirth and Gruebele, 2013), which could be consistent with macromolecular crowding at actin bundles, in line with the local diffusion slowdown we unveiled through SM*d*M.

For the nucleus, we visualized diffusion slowdown at the nanoscale due to macromolecular crowding at the nucleolus and the chromatin. Although single-location FCS measurements have previously shown reduced *D* at the chromatin and the nucleolus (Bancaud et al., 2009), FCS mapping in ~1 μm-spaced arrays finds no correlation between *D* and chromatin structure (Dross et al., 2009). Our correlated SM*d*M and SMLM results helped establish a definite association, at the nanoscale, between local *D* and the chromatin ultrastructure. The SM*d*M-resolved coexistence of fast and slow diffusion domains in the nucleus may be functionally important, as envisioned by the chromosome-territory– interchromatin-compartment (CT-IC) model (Cremer and Cremer, 2001).

Following a surprising observation we made with mEos3.2-NLS, we next unveiled an unexpected, dominating role of protein net charge on intracellular diffusion. Intriguingly, SM*d*M revealed that whereas a negative net charge did not significantly affect protein diffusion, the possession of positive net charges was a key factor for diffusion slowdown, and the degree of slowdown depended on the specific subcellular environments (Figure 4F). Interestingly, in bacteria, a recent study (Schavemaker et al., 2017) has examined the diffusion of differently charged GFP variants, and also finds that all negatively charged and neutral GFPs diffuse alike, whereas positively charged GFPs diffuse much slower, a result ascribed to interaction with ribosomes. The mammalian cell, however, is a much more complicated system.

Notably, the mammalian cytosol contains a high (~150 mM) concentration of small cations, notably K^+^, whereas the total concentration of small anions is disproportionally low (~15 mM) (Lodish et al., 2003). Charge balance thus mandates intracellular bio(macro)molecules to take the negative charges. Whereas the backbones of DNA and RNA are known to be negatively charged, the significance of protein net charges has started to gain attention in recent years (Borgia et al., 2018; Gitlin et al., 2006; Mu et al., 2017; Schavemaker et al., 2017; Smith et al., 2016). We noticed that the most abundant proteins in the mammalian cytoplasm tend to be either strongly negatively charged or neutral (Table S2). For instance, two abundant molecular chaperones, Hsp90ab1 and HspA8, carry −39 and −13 net charges, respectively. Earlier analysis not considering relative abundances also suggests a majority of cytoplasmic proteins to be negatively charged (Schwartz et al., 2001).

Consequently, the peculiar, sign-asymmetric dependency of *D* on net charge we observed (Figure 4F) may be rooted in the asymmetric intracellular abundance of positively charged, small metal ions vs. negatively charged, large biomolecules. For intracellular diffusion, whereas a negatively charged diffuser is readily neutralized by the abundant small cations and so behaves similarly as neutral diffusers (Figure 6A), a positively charged diffuser is dragged down by large biomolecules that are predominantly negatively charged (Figure 6B). Indeed, *in vitro* experiments have shown that the diffusion of charged proteins in polymeric solutions to be substantially impeded by opposite-charge, but not same-charge or neutral, polymers (Zustiak et al., 2011). At a fundamental level, such charge-asymmetric impediments to diffusion may, conversely, explain the preponderance of negatively charged proteins in the cell we noted above (Table S2): the cell may have evolved to agree on a negatively charged convention to minimize nonspecific interactions and diffusion slowdown, since DNA and RNA are already negatively charged.

**Figure 6.**
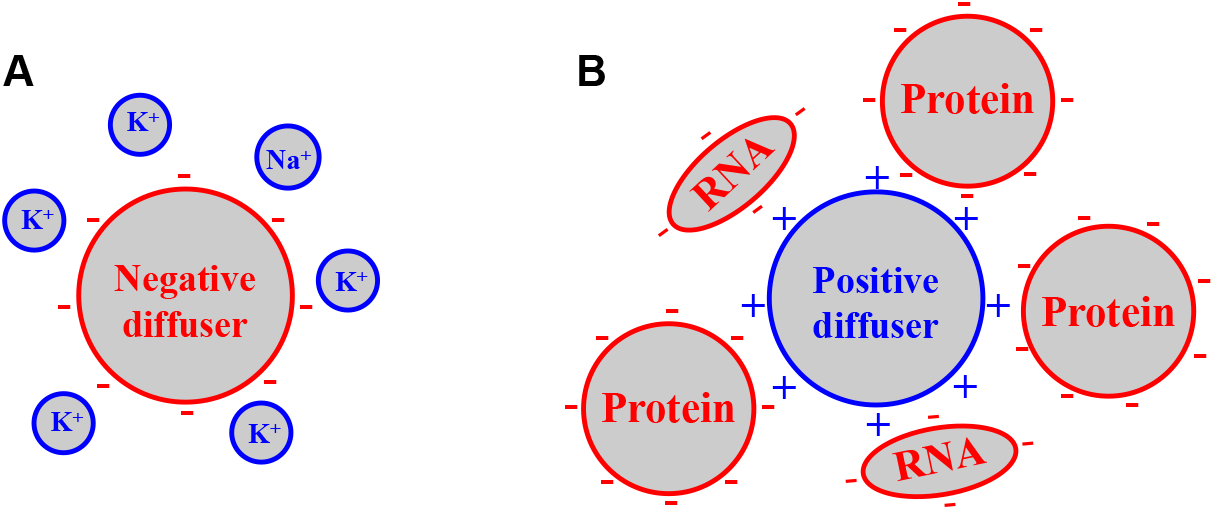
Asymmetric effects of negative and positive net charges on intracellular diffusion. (A) A negatively charged diffuser is readily neutralized by the abundant, small metal cations inside the cell, and so diffuses similarly as neutral counterparts. (B) A positively charged diffuser is not effectively neutralized/screened by the very limited amount of intracellular small anions; its dynamic interactions with the negatively charged, large biomolecules insides the cell substantially hinder diffusion.

From a different standpoint, our observed strong dependence of diffusion rate, and thus nonspecific protein interactions, on positive net charges also calls for a reexamination of previous work in which FPs or other probes may have inadvertently shifted the protein net charge. Indeed, many common FPs are highly negatively charged (*e.g.*, −7 charges for most GFP derivatives, including EGFP, ECFP, and Venus), and hence could have biased experimental results towards the negative-charge regime, where the true effects of net charge are masked (Figure 4F).

Together, we have shown how the local statistics of instantaneous displacements of unbound single molecules can unveil rich, nanoscale heterogeneities in intracellular diffusivity. Whereas fascinating results were obtained here with free FPs, we expect SM*d*M with FP-tagged proteins to be powerful in probing specific protein-protein interactions. The further integration of SM*d*M with other emerging super-resolution methods, *e.g.*, spectrally resolved SMLM (Yan et al., 2018), represents additional exciting possibilities.

## EXPERIMENTAL PROCEDURES

### Optical setup

Single-molecule experiments were performed on a Nikon Ti-E inverted fluorescence microscope. Lasers at 561 nm (OBIS 561 LS, Coherent, 165 mW, for excitation of fluorescence) and 405 nm (Stradus 405, Vortran, 100 mW, for photoactivation) were collinearly combined and focused at the back focal plane of an oil-immersion objective lens (Nikon CFI Plan Apochromat λ 100×, NA 1.45) through a dichroic mirror (ZT561rdc, Chroma). A translation stage shifted the laser beams toward the edge of the objective lens so that the light reached the sample at an incidence angle slightly smaller than the critical angle of the glass-water interface, thus illuminating a few micrometers into the sample. Fluorescence emission was filtered by a long-pass filter (ET575lp, Chroma) and an additional band-pass filter (ET605/70m, Chroma) in front of the EMCCD camera (iXon Ultra 897, Andor). Both the excitation laser (561 nm) and the photoactivation laser (405 nm) were modulated by a multifunction I/O board (PCI-6733, National Instruments), which also read the camera exposure output TTL signal for synchronization.

### Plasmid constructs

mEos3.2-C1 was a gift from Michael Davidson & Tao Xu (Addgene plasmid # 54550) (Zhang et al., 2012), and was used without modification as the “free” version of mEos3.2 (+2 net charge). The sequences of mEos3.2-NLS and the other modified constructs are listed in Table S1. mEos3.2-NLS was constructed by inserting the desired DNA sequence (Integrated DNA Technologies) between the SalI and BamHI restriction enzyme recognition sites within the short sequence at the C-terminus of mEos3.2-C1. mEOS3.2(+7b) was constructed by replacing the DNA strains after the Kpn2I restriction enzyme recognition site with the desired DNA sequence. Other mEos3.2-based versions were prepared by inserting the desired DNA sequences at the EcoRI restriction enzyme recognition site. The +7 charged mEosP5-C1(+7c) was constructed by replacing DNA strains in mEos3.2-C1 with the desired sequences between the AgeI and EcoRV restriction enzyme recognition sites and between the PflMI and Kpn2I restriction enzyme recognition sites. Verification of plasmid constructs was confirmed through Sanger sequencing. Net charges of the proteins were estimated by summing the charge of each amino acid or via the online tool Protein Calculator v3.4 (http://protcalc.sourceforge.net/), yielding comparable results (see details in the notes below Table S1).

### Cell culturing and transfection

18-mm diameter glass coverslips were cleaned with a heated piranha solution (sulfuric acid and hydrogen peroxide at 3:1), and then rinsed with Milli-Q water (18.4 MΩ cm). Ptk2 and U2OS cells were cultured in Dulbecco’s Modified Eagle’s Medium (DMEM) with 10% fetal bovine serum (FBS), 1× GlutaMAX Supplement, and 1× non-essential amino acids (NEAA) in 5% CO_2_ at 37 ^°^C. 24 hours before imaging, cells were transfected with the Neon Transfection System (ThermoFisher) according to the recommended protocol, and then plated onto the pre-cleaned glass coverslips at a density of ~40,000/cm^2^.

### SMdM of live cells

SM*d*M of live cells was performed in a Leibovitz’s L-15 medium containing 20 mM HEPES buffer, except for the hyperosmotic experiment, for which additional glucose was added at 49 mg/mL. For a typical recorded frame size of 256×256 pixels (~41×41 μm^2^ sample area), the EMCCD camera exposure time and dead time were 9.0 ms and 157 μs, respectively, hence a frame rate of 109.3 frames per second. To access sub-frame temporal resolution, for each paired frames, two excitation (561 nm) pulses of duration *τ* (500 μs typical) were placed towards the end of the first frame and the beginning of the second frame, respectively (Figure 1C), at a center-to-center separation of Δ*t* (1 ms typical, but 5 ms for the very slow diffusion in the NLS and +14 charged samples). The wait time between the two excitation pulses was evenly distributed across the EMCCD dead time. For photoactivation, a low level of 405 nm laser was applied during the first half of the first frame in each paired frames to achieve a low density of emitting single molecules across the view. In a typical experiment, the estimated peak and average power densities of the 561 nm excitation laser at the sample were ~6 and 0.3 kW/cm^2^, respectively. The average power density of the 405 nm activation laser was usually 0-0.05 W/cm^2^, depending on the expression level. The above scheme of photoactivation and paired excitation was repeated many times (5-7×10^4^ typical) to generate the final SM*d*M data.

### SMLM imaging of fixed cells after live-cell SMdM

After the above SM*d*M experiment on live cells, the sample was chemically fixed for subsequent fluorescent labeling and SMLM imaging. For SMLM of the actin cytoskeleton, the cells were fixed with 0.3% glutaraldehyde and 0.25% Triton X-100 in the cytoskeleton buffer (10 mM MES [2-(N-morpholino)ethanesulfonic acid] buffer, 150 mM NaCl, 5 mM EGTA (ethylene glycol tetraacetic acid), 5 mM glucose, 5 mM MgCl_2_, pH 6.1) for 1 minute, then fixed with 2% glutaraldehyde in the cytoskeleton buffer for 30 minutes (Xu et al., 2012). The sample was then treated with a 0.1% NaBH_4_ solution in phosphate-buffered saline (PBS) for 5 minutes × 2 times, and then washed with PBS for 10 minutes for 3 times. Actin was labeled with 0.5 μM Alexa Fluor 647-phalloidin (Invitrogen A22287) solution in PBS for 30 minutes, and then washed with PBS for 5 minutes × 2 times. For SMLM of DNA, the cells were fixed with 4% paraformaldehyde in PBS and washed with PBS for 10 minutes × 3 times. Then the DNA was labeled with NucSpot Live 650 (Biotium #40082) in PBS (1:1000) for 20 minutes. The sample was washed with PBS for 5 minutes × 2 times. SMLM was performed on the same microscope setup using a 642 nm laser (Stradus 642, Vortran, 110 mW). The SMLM imaging buffer was PBS containing 5% glucose, 200 mM cysteamine, 0.8 mg/mL glucose oxidase, and 40 μg/mL catalase. The acquired SMLM data were processed as described previously (Rust et al., 2006).

### Data analysis for SMdM

Single-molecule images were first super-localized as described previously (Rust et al., 2006). For each pair of frames, the super-localized positions of the molecules identified in the second frame were used to search for matching molecules in the first frame within a cutoff radius *R* (800 nm typical). Displacements (*d*) were calculated for the matched molecules, and the process was repeated for all the paired frames. The resultant, accumulated *d* values were spatially binned onto 100×100 nm^2^ grids for Figures 1-2, and 120×120 nm^2^ grids for Figures 3-5 and S1-S3. The distribution of *d* in each spatial bin was next individually fitted through maximum likelihood estimation (MLE) to determine local *D*. The extraction of *D* from the distribution of single-step displacement has been previously examined (Anderson et al., 1992; Hansen et al., 2018; Kues et al., 2001; Lin et al., 2014), typically using frame-to-frame displacements from long trajectories of individual particles. In SM*d*M, fitting is instead for different molecules that visit a given location for just a pair of frames in the very short duration of Δ*t*, and we add one more term to accommodate mismatched molecules. According to two-dimensional random walk (since in our measurements we do not measure the axial position and only calculate the in-plane displacement), the probability density for a particle to move a distance *r* in the fixed time interval Δ*t* is (Anderson et al., 1992; Kues et al., 2001; Lin et al., 2014):

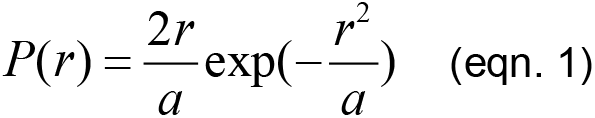

where *a* = 4*D*Δ*t*. Assuming the density of background molecules (mismatches in pairing) to be spatially homogeneous within the search radius, the probability of finding a background molecule between *r* and *r*+d*r* is proportional to the area 2*πr*d*r*, which increases linearly with *r*. We thus modified eqn. 1 to account for this background effect:

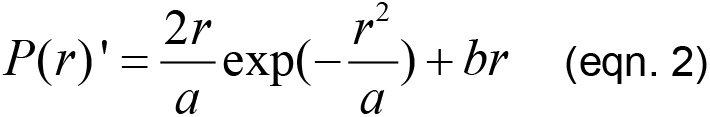

where *b* fits to the slope of a linearly increasing background. Using Eqn. 2 to fit the SM*d*M data through MLE yielded robust results for experiments carried out at different single-molecule densities (Figure S1).

## SUPPLEMENTAL INFORMATION

Supplemental Information includes three figures and two tables.

## ACKNOWLEDGMENTS

We thank Seonah Moon for discussion, and Manni He and Yennie Shyu for help with preparation of the DNA constructs. This work was supported by the National Institute Of General Medical Sciences of the National Institutes of Health (DP2GM132681), the Beckman Young Investigator Program, and the Packard Fellowships for Science and Engineering. K.X. is a Chan Zuckerberg Biohub investigator.

## AUTHOR CONTRIBUTIONS

K. X. conceived the research. L. X. and K. C. designed and conducted the experiments. All authors contributed to experimental designs, data analysis, and paper writing.

## REFERENCES

Anderson, C.M., Georgiou, G.N., Morrison, I.E.G., Stevenson, G.V.W., and Cherry, R.J. (1992). Tracking of cell surface receptors by fluorescence digital imaging microscopy using a charge-coupled device camera. Low-density lipoprotein and influenza virus receptor mobility at 4 °C. J Cell Sci 101, 415–425.

Bancaud, A., Huet, S., Daigle, N., Mozziconacci, J., Beaudouin, J., and Ellenberg, J. (2009). Molecular crowding affects diffusion and binding of nuclear proteins in heterochromatin and reveals the fractal organization of chromatin. EMBO J 28, 3785–3798.

Baum, M., Erdel, F., Wachsmuth, M., and Rippe, K. (2014). Retrieving the intracellular topology from multi-scale protein mobility mapping in living cells. Nat Commun 5, 4494.

Benke, A., and Manley, S. (2012). Live-cell dSTORM of cellular DNA based on direct DNA labeling. ChemBioChem 13, 298–301.

Betzig, E., Patterson, G.H., Sougrat, R., Lindwasser, O.W., Olenych, S., Bonifacino, J.S., Davidson, M.W., Lippincott-Schwartz, J., and Hess, H.F. (2006). Imaging intracellular fluorescent proteins at nanometer resolution. Science 313, 1642–1645.

Boersma, A.J., Zuhorn, I.S., and Poolman, B. (2015). A sensor for quantification of macromolecular crowding in living cells. Nat Methods 12, 227–229.

Boisvert, F.M., van Koningsbruggen, S., Navascues, J., and Lamond, A.I. (2007). The multifunctional nucleolus. Nat Rev Mol Cell Biol 8, 574–585.

Borgia, A., Borgia, M.B., Bugge, K., Kissling, V.M., Heidarsson, P.O., Fernandes, C.B., Sottini, A., Soranno, A., Buholzer, K.J., Nettels, D., et al. (2018). Extreme disorder in an ultrahigh-affinity protein complex. Nature 555, 61–66.

Cognet, L., Leduc, C., and Lounis, B. (2014). Advances in live-cell single-particle tracking and dynamic super-resolution imaging. Curr Opin Chem Biol 20, 78–85.

Cremer, T., and Cremer, C. (2001). Chromosome territories, nuclear architecture and gene regulation in mammalian cells. Nat Rev Genet 2, 292–301.

Digman, M.A., and Gratton, E. (2011). Lessons in fluctuation correlation spectroscopy. Annu Rev Phys Chem 62, 645–668.

Dross, N., Spriet, C., Zwerger, M., Muller, G., Waldeck, W., and Langowski, J. (2009). Mapping eGFP oligomer mobility in living cell nuclei. PLoS One 4, e5041.

Ebbinghaus, S., Dhar, A., McDonald, D., and Gruebele, M. (2010). Protein folding stability and dynamics imaged in a living cell. Nat Methods 7, 319–323.

Elf, J., and Barkefors, I. (2019). Single-molecule kinetics in living cells. Annu Rev Biochem 88, 635–659.

Elf, J., Li, G.W., and Xie, X.S. (2007). Probing transcription factor dynamics at the single-molecule level in a living cell. Science 316, 1191–1194.

Enderlein, J., Gregor, I., Patra, D., Dertinger, T., and Kaupp, U.B. (2005). Performance of fluorescence correlation spectroscopy for measuring diffusion and concentration. ChemPhysChem 6, 2324–2336.

English, B.P., Hauryliuk, V., Sanamrad, A., Tankov, S., Dekker, N.H., and Elf, J. (2011). Single-molecule investigations of the stringent response machinery in living bacterial cells. Proc Natl Acad Sci U S A 108, E365–E373.

Gitlin, I., Carbeck, J.D., and Whitesides, G.M. (2006). Why are proteins charged? Networks of charge-charge interactions in proteins measured by charge ladders and capillary electrophoresis. Angew Chem-Int Edit 45, 3022–3060.

Hansen, A.S., Woringer, M., Grimm, J.B., Lavis, L.D., Tjian, R., and Darzacq, X. (2018). Robust model-based analysis of single-particle tracking experiments with Spot-On. eLife 7, e33125.

Hess, S.T., Girirajan, T.P.K., and Mason, M.D. (2006). Ultra-high resolution imaging by fluorescence photoactivation localization microscopy. Biophys J 91, 4258–4272.

Ishikawa-Ankerhold, H.C., Ankerhold, R., and Drummen, G.P.C. (2012). Advanced fluorescence microscopy techniques-FRAP, FLIP, FLAP, FRET and FLIM. Molecules 17, 4047–4132.

Kues, T., Peters, R., and Kubitscheck, U. (2001). Visualization and tracking of single protein molecules in the cell nucleus. Biophys J 80, 2954–2967.

Kuimova, M.K., Botchway, S.W., Parker, A.W., Balaz, M., Collins, H.A., Anderson, H.L., Suhling, K., and Ogilby, P.R. (2009). Imaging intracellular viscosity of a single cell during photoinduced cell death. Nat Chem 1, 69–73.

Kusumi, A., Tsunoyama, T.A., Hirosawa, K.M., Kasai, R.S., and Fujiwara, T.K. (2014). Tracking single molecules at work in living cells. Nat Chem Biol 10, 524–532.

Lin, W.C., Iversen, L., Tu, H.L., Rhodes, C., Christensen, S.M., Iwig, J.S., Hansen, S.D., Huang, W.Y.C., and Groves, J.T. (2014). H-Ras forms dimers on membrane surfaces via a protein-protein interface. Proc Natl Acad Sci U S A 111, 2996–3001.

Lippincott-Schwartz, J., Snapp, E., and Kenworthy, A. (2001). Studying protein dynamics in living cells. Nat Rev Mol Cell Biol 2, 444–456.

Lodish, H., Berk, A., Matsudaira, P., Kaiser, C.A., Krieger, M., Scott, M.P., Zipursky, L., and Darnell, J. (2003). In Molecular Cell Biology (New York: W.H. Freeman), p. 253.

Machan, R., and Wohland, T. (2014). Recent applications of fluorescence correlation spectroscopy in live systems. FEBS Lett 588, 3571–3584.

Manley, S., Gillette, J.M., Patterson, G.H., Shroff, H., Hess, H.F., Betzig, E., and Lippincott-Schwartz, J. (2008). High-density mapping of single-molecule trajectories with photoactivated localization microscopy. Nat Methods 5, 155–157.

Manzo, C., and Garcia-Parajo, M.F. (2015). A review of progress in single particle tracking: from methods to biophysical insights. Rep Prog Phys 78, 124601.

Marfori, M., Mynott, A., Ellis, J.J., Mehdi, A.M., Saunders, N.F.W., Curmi, P.M., Forwood, J.K., Boden, M., and Kobe, B. (2011). Molecular basis for specificity of nuclear import and prediction of nuclear localization. Biochim Biophys Acta-Mol Cell Res 1813, 1562–1577.

Milo, R., and Phillips, R. (2016). Cell Biology by the Numbers (New York, NY: Garland Science).

Mu, X., Choi, S., Lang, L., Mowray, D., Dokholyan, N.V., Danielsson, J., and Oliveberg, M. (2017). Physicochemical code for quinary protein interactions in *Escherichia coli*. Proc Natl Acad Sci U S A 114, E4556–E4563.

Potma, E.O., de Boeij, W.P., Bosgraaf, L., Roelofs, J., van Haastert, P.J.M., and Wiersma, D.A. (2001). Reduced protein diffusion rate by cytoskeleton in vegetative and polarized *Dictyostelium* cells. Biophys J 81, 2010–2019.

Ricci, M.A., Manzo, C., Garcia-Parajo, M.F., Lakadamyali, M., and Cosma, M.P. (2015). Chromatin fibers are formed by heterogeneous groups of nucleosomes *in vivo*. Cell 160, 1145–1158.

Ries, J., and Schwille, P. (2012). Fluorescence correlation spectroscopy. Bioessays 34, 361–368.

Rivas, G., and Minton, A.P. (2016). Macromolecular crowding *in vitro*, *in vivo*, and in between. Trends Biochem Sci 41, 970–981.

Rust, M.J., Bates, M., and Zhuang, X. (2006). Sub-diffraction-limit imaging by stochastic optical reconstruction microscopy (STORM). Nat Methods 3, 793–795.

Schavemaker, P.E., Smigiel, W.M., and Poolman, B. (2017). Ribosome surface properties may impose limits on the nature of the cytoplasmic proteome. eLife 6, e30084.

Schwartz, R., Ting, C.S., and King, J. (2001). Whole proteome pI values correlate with subcellular localizations of proteins for organisms within the three domains of life. Genome Res 11, 703–709.

Seksek, O., Biwersi, J., and Verkman, A.S. (1997). Translational diffusion of macromolecule-sized solutes in cytoplasm and nucleus. J Cell Biol 138, 131–142.

Smith, A.E., Zhou, L.Z., Gorensek, A.H., Senske, M., and Pielak, G.J. (2016). In-cell thermodynamics and a new role for protein surfaces. Proc Natl Acad Sci U S A 113, 1725–1730.

Swaminathan, R., Hoang, C.P., and Verkman, A.S. (1997). Photobleaching recovery and anisotropy decay of green fluorescent protein GFP-S65T in solution and cells: Cytoplasmic viscosity probed by green fluorescent protein translational and rotational diffusion. Biophys J 72, 1900–1907.

Szczurek, A., Klewes, L., Xing, J., Gourram, A., Birk, U., Knecht, H., Dobrucki, J.W., Mai, S., and Cremer, C. (2017). Imaging chromatin nanostructure with binding-activated localization microscopy based on DNA structure fluctuations. Nucleic Acids Res 45, e56.

Wirth, A.J., and Gruebele, M. (2013). Quinary protein structure and the consequences of crowding in living cells: Leaving the test-tube behind. Bioessays 35, 984–993.

Xiong, K., and Blainey, P.C. (2016). Molecular sled sequences are common in mammalian proteins. Nucleic Acids Res 44, 2266–2273.

Xu, K., Babcock, H.P., and Zhuang, X. (2012). Dual-objective STORM reveals three-dimensional filament organization in the actin cytoskeleton. Nature Methods 9, 185–188.

Yan, R., Moon, S., Kenny, S.J., and Xu, K. (2018). Spectrally resolved and functional super-resolution microscopy via ultrahigh-throughput single-molecule spectroscopy. Acc Chem Res 51, 697–705.

Yang, Z.G., Cao, J.F., He, Y.X., Yang, J.H., Kim, T., Peng, X.J., and Kim, J.S. (2014). Macro-/micro-environment-sensitive chemosensing and biological imaging. Chem Soc Rev 43, 4563–4601.

Zhang, M.S., Chang, H., Zhang, Y.D., Yu, J.W., Wu, L.J., Ji, W., Chen, J.J., Liu, B., Lu, J.Z., Liu, Y.F., et al. (2012). Rational design of true monomeric and bright photoactivatable fluorescent proteins. Nat Methods 9, 727–729.

Zustiak, S.P., Nossal, R., and Sackett, D.L. (2011). Hindered diffusion in polymeric solutions studied by fluorescence correlation spectroscopy. Biophys J 101, 255–264.

