## SUPPLEMENTAL INFORMATION for "Single-molecule displacement mapping unveils nanoscale heterogeneities and charge effects in intracellular diffusivity"

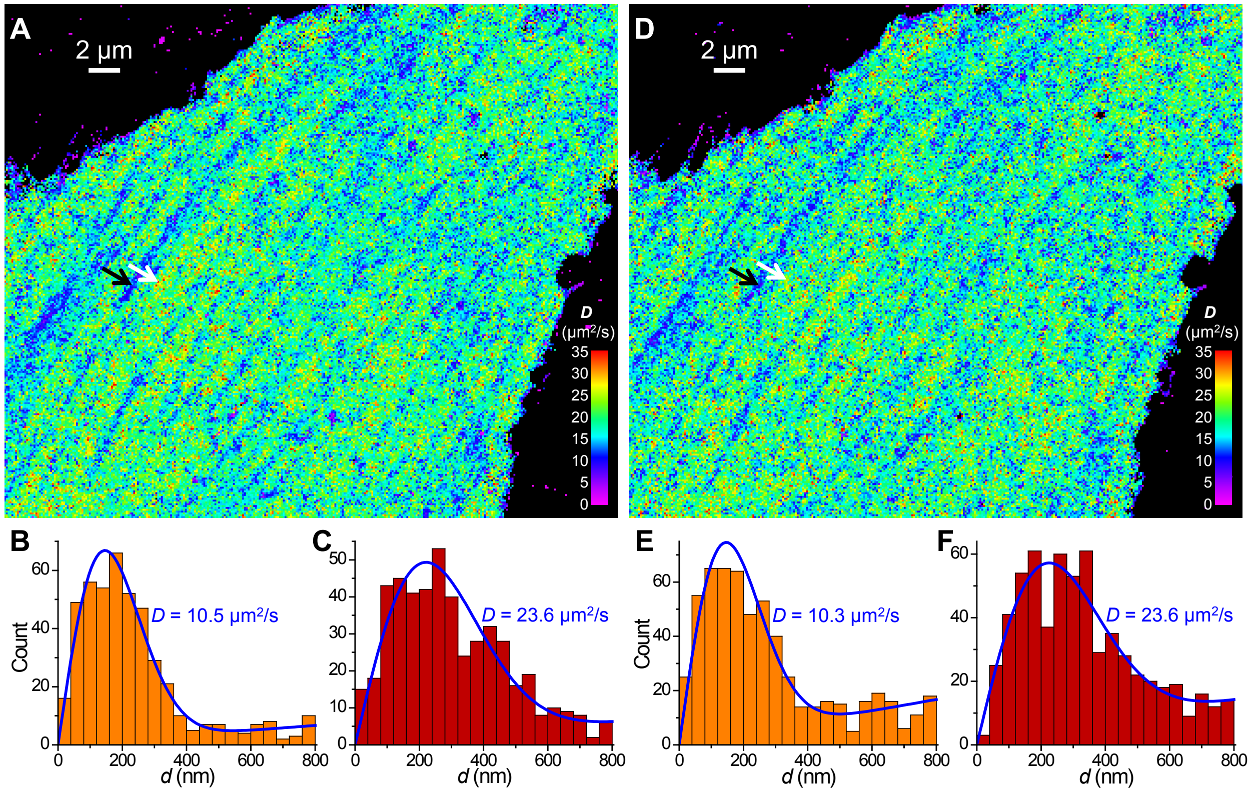


**Figure S1. SM*d*M results at different single-molecule densities, related to Figure 1**

Free mEos3.2 was expressed in the cytoplasm of a PtK2 cell, and SM*d*M was performed on the same cell at a low single-molecule density for 60,000 pairs of pulses (A-C), or at a high single-molecule density for 30,000 pairs of pulses (D-F) by increasing the power of the photoactivation (405 nm) laser.

(A) SM*d*M diffusivity map for the low single-molecule density experiment, obtained by spatially binning the single-molecule displacement *d* data onto 120×120 $nm^2$ grids, and then individually fitting the distribution of *d* in each bin to eqn. 2 through MLE.

(B,C) Distribution of *d* for two 360×360 $nm^2$ areas pointed to by the black (for B) and white (for C) arrows in (A), respectively. Blue lines are MLE results using eqn. 2, with the resultant *D* values labeled in each figure.

(D-F) Results of the high single-molecule density experiment: comparable *D* values are obtained with the much-reduced number of pulse pairs, despite an increased background due to single-molecule mismatch.

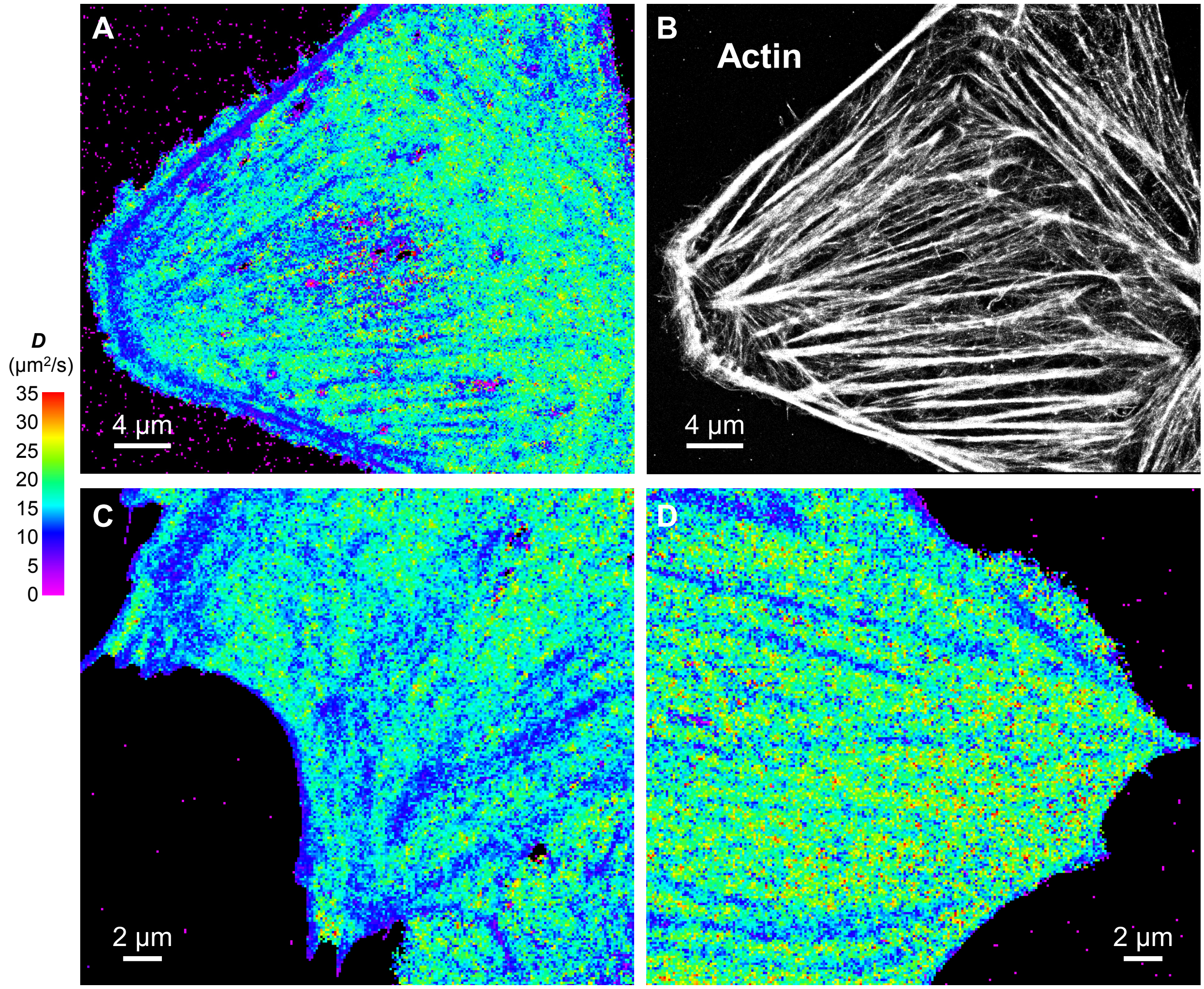


**Figure S2. Additional SM*d*M results of free mEos3.2 in the cytoplasm of live U2OS and PtK2 cells, related to Figure 2**

(A,B) Correlated SM*d*M diffusivity map for a live U2OS cell (A) vs. SMLM image of Alexa Fluor 647 phalloidin-labeled actin in the fixed cell (B).

(C,D) Additional SM*d*M diffusivity maps for the cytoplasm of PtK2 cells.

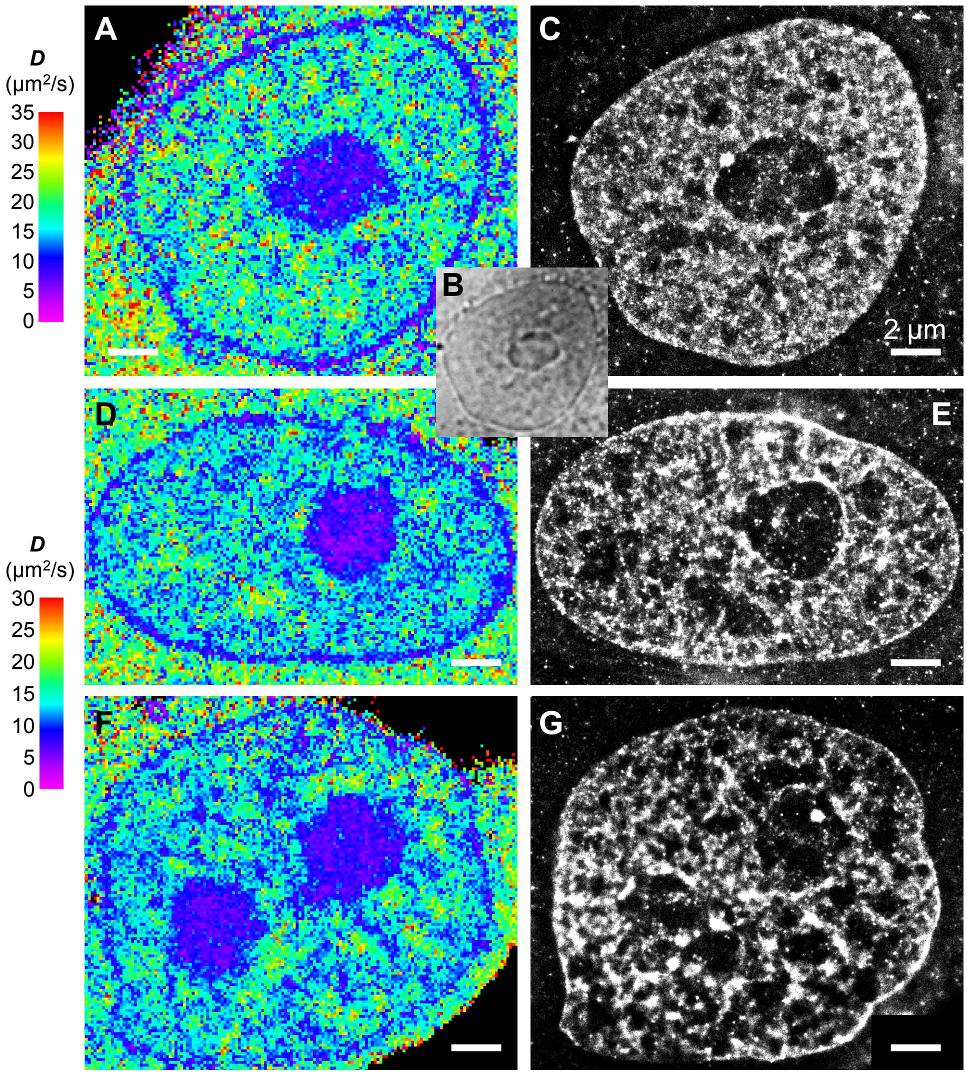


**Figure S3. Additional SM*d*M results of free mEos3.2 in the nuclei of live PtK2 cells, related to Figure 3**

(A,D,F) SM*d*M diffusivity maps of 3 different cells.

(B) Bright-field transmission image of the same view as (A), visualizing the nucleolus.

(C,E,G) SMLM images of the fixed cells in (A,D,F) using the DNA stain NucSpot Live 650. We note that as the SMLM of DNA was performed after fixation and multiple washing steps, it was difficult to image at exactly the same focal plane as the live-cell SM*d*M experiment, which accounts for some of the apparent structural mismatches.

Scale bars in all panels: 2 μm.

**Table S1**. List of plasmid constructs used in this work. Blue and red colors mark positively and negatively charged AAs that are varied between the different constructs.

| Plasmid | Protein sequence | Size (AA) | Net charge[a] | Net charge[b] |
|---|---|---|---|---|
| mEos3.2 -C1 | mEos3.2-SGLRSRAQASNSAVDGTAGPGSTGSR<br>(mEos3.2 = MSAIKPDMKIKLRMEGNVNGHHFVIDGDGTGKPFEGKQSMDLEVKEGGPLPFAFDILTTAFHYGNRVFAKYPDNIQDYFKQSFPKGYSWERSLTFEDGGICNARNDITMEGDTFYNKVRFYGTNFPANGPVMQKKTLKWEPSTEKMYVRDGVLTGDIEMALLLEGNAHYRCDFRTTYKAKEKGVKLPGAHFVDHCIEILSHDKDYNKVKLYEHAVAHSGLPDNARR) | 252 | +2.2 | +2.5 |
| mEos3.2 -NLS | mEos3.2-SGLRSRADPKKKRKVDPKKKRKVDPKKKRKVGSTGSR | 262 | +15.2 | +15.5 |
| mEos3.2 (−14) | mEos3.2-SGLRSRAQASNSDEDEEDDEEDEEDDEDNSAVDGTAGPGSTGSR | 270 | −13.8 | −13.4 |
| mEos3.2 (−7) | mEos3.2-SGLRSRAQASNSDEDEEDDEENSAVDGTAGPGSTGSR | 263 | −6.8 | −6.5 |
| mEos3.2 (0) | mEos3.2-SGLRSRAQASNSDESTQNSAVDGTAGPGSTGSR | 259 | +0.2 | +0.5 |
| mEos3.2 (+7a) | mEos3.2-SGLRSRAQASNSKKRKRNSAVDGTAGPGSTGSR | 259 | +7.2 | +7.5 |
| mEos3.2 (+14) | mEos3.2-SGLRSRAQASNSKKRKKRKKRKKRNSAVDGTAGPGSTGSR | 266 | +14.2 | +14.5 |
| mEos3.2 (+7b) | mEos3.2- SGRQKGHKCIRLPKVNQRMSR | 247 | +7.2 | +7.5 |
| mEosP5-C1 (+7c) | mEosP5-SGLRSRAQASNSAVDGTAGPGSTGSR<br>(mEosP5 = MKSAIKPDMKIKLRMEGNVNGHHFVIDGDGTGKPFEGKQSMDLEVKKGGPLPFAFDILTTAFHYGNRVFAKYPDNIQDYFKQSFPKGYSWERSLTFEDGGICNARNDITMEGDTFYNKVRFYGTNFPANGPVMQKKTLKWEPSTEKMYVRDGVLTGDIEMALLLEGNAHYRCDFRTTYKAKEKGVKLPGAHFVDHCIEILSHDKKYNKVKLYEHAVAHSGLPDNARR) | 253 | +7.2 | +7.5 |

a: Calculated by summing the charges of each amino acid at pH 7.4 (Requiao et al., 2017): lysine = +0.999; arginine = +1.000; histidine = +0.048; glutamic acid = −0.999; aspartic acid = −1.000; cysteine = −0.085, and all other amino acids = 0.000.

b: Calculated via the online tool Protein Calculator v3.4 (http://protcalc.sourceforge.net/) for pH 7.4.

**Table S2.** List of estimated net charges for the most abundant (>0.2% of total protein mass) cytoplasmic proteins, based on the proteomics of the U2OS human cell line (Beck et al., 2011; Liebermeister et al., 2014). Protein sequences are from UniProt (https://www.uniprot.org/). Net charges are estimated for pH 7.4 via Protein Calculator (http://protcalc.sourceforge.net/). For each category, proteins are listed in the order of mass abundance (% of the total protein mass of the cell). This showed that most proteins in the categories of "cytoskeletal proteins", "chaperones and folding catalysts", and "others" are either strongly negatively charged (<−10) or neutral (within ±2). Half of the proteins in the "glycolysis" group are mildly (~+3) positive, possibly for their intended interactions with the negatively charged, phosphorylated glucose metabolites, which thereby neutralizing the total charge. Three proteins in the group of "ribosome" and one in "translation factors" are strongly positively charged, but these positive charges are more than compensated by their binding partner, the heavily negatively charged RNA (Knight et al., 2013; Schavemaker et al., 2017).

**Cytoskeletal proteins & regulators**

| Name | %mass | Net charge |
|---|---|---|
| Vim | 2.7 | -18.6 |
| TubA1c | 2.2 | -22.7 |
| ActB | 2.2 | -11.7 |
| Cfl1 | 1.2 | 1.6 |
| FlnA | 0.84 | -50.8 |
| Plec | 0.83 | -76.9 |
| myh9 | 0.71 | -45.9 |
| pfn1 | 0.68 | 1.8 |
| FlnB | 0.36 | -60.9 |
| FlnC | 0.32 | -54.6 |
| TubB6 | 0.30 | -24.7 |
| LmnA | 0.29 | -2.1 |
| Myl6 | 0.28 | -14.0 |
| SptAn1 | 0.27 | -106.4 |

**Chaperones and folding catalysts**

| Name | %mass | Net charge |
|---|---|---|
| Hsp90ab1 | 2.2 | -39.3 |
| HspA8 | 2.0 | -12.8 |
| cct2 | 1.1 | -7.9 |
| PPIA | 0.97 | 0.9 |
| HspD1 | 0.89 | -5.2 |
| HspB1 | 0.60 | -2.7 |
| cct6a | 0.40 | -4.5 |
| cct5 | 0.20 | -13.6 |

**Glycolysis**

| Name | %mass | Net charge |
|---|---|---|
| Pkm2 | 3.1 | 2.4 |
| Eno1 | 2.5 | 0.0 |
| GAPDH | 2.0 | 3.6 |
| Tpi1 | 1.7 | -5.0 |
| AldOa | 1.1 | 3.2 |
| Pgk1 | 0.85 | 2.8 |
| LdhA | 0.53 | 3.2 |
| Eno3 | 0.48 | 1.1 |
| Eno2 | 0.22 | -18.0 |

**Ribosome**

| Name | %mass | Net charge |
|---|---|---|
| Rpl37a | 0.54 | 18.7 |
| Rps15a | 0.26 | 10.1 |
| Rpl7a | 0.25 | 40.9 |
| RplP0 | 0.20 | -4.0 |

**Translation factors**

| Name | %mass | Net charge |
|---|---|---|
| Eef1a1 | 2.7 | 11.4 |
| Eef2 | 1.1 | -4.8 |
| Eif5a | 0.50 | -7.1 |
| Eef1d | 0.39 | -14.8 |

**Others**

| Name | %mass | Net charge |
|---|---|---|
| Mif | 1.9 | 0.8 |
| LgaLs1 | 0.79 | -3.4 |
| Tkt | 0.47 | 1.7 |
| Cltc | 0.38 | -39.8 |
| GstP1 | 0.33 | -3.3 |
| Eif4a1 | 0.26 | -9.0 |
| FasN | 0.23 | -34.1 |

### REFERENCES FOR SUPPLEMENTAL INFORMATION

Beck, M., Schmidt, A., Malmstroem, J., Claassen, M., Ori, A., Szymborska, A., Herzog, F., Rinner, O., Ellenberg, J., and Aebersold, R. (2011). The quantitative proteome of a human cell line. Mol Syst Biol *7*, 549.

Knight, A.M., Culviner, P.H., Kurt-Yilmaz, N., Zou, T.S., Ozkan, S.B., and Cavagnero, S. (2013). Electrostatic effect of the ribosomal surface on nascent polypeptide dynamics. ACS Chem Biol *8*, 1195-1204.

Liebermeister, W., Noor, E., Flamholz, A., Davidi, D., Bernhardt, J., and Milo, R. (2014). Visual account of protein investment in cellular functions. Proc Natl Acad Sci U S A *111*, 8488-8493.

Requiao, R.D., Fernandes, L., de Souza, H.J.A., Rossetto, S., Domitrovic, T., and Palhano, F.L. (2017). Protein charge distribution in proteomes and its impact on translation. PLoS Comput Biol *13*, e1005549.

Schavemaker, P.E., Smigiel, W.M., and Poolman, B. (2017). Ribosome surface properties may impose limits on the nature of the cytoplasmic proteome. eLife *6*, e30084.
